# Bark beetle protein elicitors trigger biphasic immune responses in Norway spruce seedlings

**DOI:** 10.64898/2026.05.22.727111

**Authors:** Marcelo Ramires, Sigrid Netherer, Martin Schebeck, Reinhard Ertl, Muhammad Ahmad, Erwann Arc, Marcela van Loo, Carlos Trujillo-Moya

**Author notes:** Corresponding author Carlos Trujillo-Moya /. Marcelo Ramires/ Sigrid Netherer / Martin Schebeck / Reinhard Ertl / Muhammad Ahmad / Erwann Arc / Marcela van Loo /.

## Abstract

Norway spruce (*Picea abies*) responds to attacks by the spruce bark beetle (*Ips typographus*) through the rapid activation of local defense mechanisms, but field studies can be difficult to standardize due to variable attack pressure and environmental heterogeneity. Here, we developed a phytotron-based assay that mimics early beetle-associated stress using insect-derived protein extracts, enabling reproducible molecular analyses under controlled conditions. Ten-week-old spruce seedlings were stem-treated with mock buffer or beetle protein extracts, followed by transcriptomic analyses of stem tissues and targeted metabolomic profiling of needles at 2 and 48 h post-inoculation. RT-qPCR analysis revealed rapid transcriptional activation of signaling and defense genes in Norway spruce, with NP-40-based protein extracts producing the most consistent early response. RNA-seq analysis revealed transcriptional dynamics, with 488 differentially expressed genes detected at 2 h and 84 at 48 h post-inoculation relative to mock-treated controls. Early responses at 2 h were characterized by activation of genes associated with immune perception and signal transduction. By 48 h, the response shifted toward accumulation of transcripts encoding defense proteins such as chitinases, defensins, proteinase inhibitors, and pathogenesis-related (PR) proteins. Importantly, a substantial proportion of differentially expressed genes overlapped with those previously identified in mature Norway spruce trees during pioneer bark beetle attack under field conditions, supporting the biological relevance of the assay. In contrast, targeted analyses of secondary metabolites performed in needle tissue revealed limited systemic changes across time points, suggesting that early induced defenses may remain largely localized to the stem. Together, these results demonstrate that beetle-derived proteins trigger a rapid and temporally structured defense response in Norway spruce seedlings and establish a reproducible elicitor-based platform that may provide a broadly applicable framework for investigating bark beetle-induced defenses across conifers.

**Highlight:** Bark beetle protein elicitors trigger temporally structured immune responses in Norway spruce seedlings that overlap with responses observed in mature trees, with rapid immune signaling at 2 h followed by defense protein accumulation at 48 h.

## Introduction

Norway spruce (*Picea abies* [L.] H. Karst), a key species in European forests, is increasingly threatened by bark beetle infestations, particularly by the spruce bark beetle *Ips typographus*. These outbreaks, often triggered by climate-related stressors such as drought and storm, have increased in frequency and intensity, leading to large-scale forest dieback and significant economic and ecological losses (Allen et al., 2010; Hlásny et al., 2021; Seidl et al., 2015). *Picea abies* not only holds major economic value for timber production but also plays a central role in regulating hydrology, stabilizing soil, and maintaining biodiversity in alpine and subalpine ecosystems (Chakraborty et al., 2021; “The State of the World’s Forests 2020”).

*Ips typographus* is a phloem-feeding insect that colonizes mature trees by boring through the outer bark to access the nutrient-rich phloem. Once male pioneer beetles initiate an attack, mass colonization by males and females is mediated by aggregation pheromones, overwhelming the tree’s defense capacity. However, during the early stages of attack, and before colonization and widespread tissue damage, Norway spruce can mount a localized defense response that may prevent the brood establishment (Franceschi et al., 2005; Krokene, 2015). To counteract such biotic threats, conifers rely on a complex defense arsenal involving both constitutive and inducible mechanisms, including resin production, specialized secondary metabolites and defense proteins following rapid transcriptional reprogramming mediated by phytohormone signaling (Franceschi et al., 2005; Hammerbacher et al., 2019; Krokene, 2015). These responses are known to be highly localized and dynamic, particularly during the initial stages of bark beetle attack, when the tree’s ability to detect and respond rapidly is critical for its survival (Ramires et al., 2025). This early detection may involve recognition of herbivore-associated molecular patterns (HAMPs), inducing pattern-triggered immunity (PTI)-like signaling cascades that initiate downstream defense responses (Ramires et al., 2025).

The long-standing co-evolutionary history between *I. typographus* and its conifer hosts likely shaped highly specialized defense pathways (Cognato, 2015). Although defense responses of trees to bark beetle attacks have traditionally been studied under field conditions, investigating natural outbreaks poses major technical and logistical limitations. The unpredictability of infestation pressure and timing, together with variability in environmental conditions and tree physiological status, complicates data interpretation and limits reproducibility. Although methyl jasmonate treatment and inoculation with bark beetle-associated fungi are commonly used to mimic aspects of beetle attack (Erbilgin et al., 2006; Wilkinson et al., 2022; Zhao et al., 2010), tools that directly simulate insect-derived biotic stress under fully controlled conditions remain limited.

To overcome these limitations, we aimed to establish a reproducible and scalable laboratory-based assay that mimics the early stages of bark beetle colonization. Unlike existing methods that rely on invasive approaches such as mechanical wounding or inoculations with beetles or fungi, we established a highly effective setting by isolating insect-derived proteins to directly trigger plant responses. For this, we prepared protein extracts from adult male *I. typographus* and applied them to the stems of 10-week-old *P. abies* seedlings to simulate insect-associated biotic stress under controlled conditions. While juvenile seedlings are not natural hosts for bark beetle colonization, they represent a reproducible experimental model for investigating early defense signaling in Norway spruce. Following exposure, stem tissues were harvested for RNA extraction, RT-qPCR and subsequent high-throughput RNA sequencing (RNA-seq) to characterize the plant’s transcriptomic responses. The needle tissue was additionally assessed for systemic responses using targeted analyses of secondary metabolites. This study provides a proof of concept for a standardized elicitor-based method that can be applied under controlled laboratory conditions to investigate induced defense responses in Norway spruce seedlings at the molecular level. Importantly, the significant overlap between differentially expressed genes identified here and those previously detected during pioneer bark beetle attacks in mature trees supports the biological relevance of this seedling-based system. By reducing reliance on field experiments, this approach enables the molecular dissection of spruce defense responses and provides a practical platform for downstream applications such as screening resistant genotypes. Insights obtained in this system may therefore inform comparative studies of bark beetle-induced defenses across economically and ecologically important conifers.

## Materials and methods

### Bark beetle protein extraction

Bark beetles used in this study were sourced from a permanent rearing facility (BOKU University, Vienna). Insects were reared on spruce logs (approximately 60 cm in length and 20 cm in diameter) under controlled environmental conditions (25 °C, 16:8 h light:dark photoperiod). Freshly emerged adult males, which had completed their maturation feeding, were collected 1-2 days prior to the experiments and immediately snap-frozen in liquid nitrogen. Only male beetles were selected, as they initiate attacks under natural conditions. Sex determination was performed following the morphological criteria described by Schlyter & Cederholm (1981).

Frozen individuals were ground to a fine powder using a pre-chilled mortar and pestle under liquid nitrogen. Proteins were extracted using two lysis buffers containing 50 mM Tris-HCl (pH 7.5), 150 mM NaCl, and either 1% CHAPS (a zwitterionic detergent) or 1% NP-40 (a non-ionic detergent), both supplemented with an EDTA-free protease inhibitor cocktail (Roche GmbH, Vienna, Austria). One milliliter of buffer was added per 10 beetles. The homogenates were incubated overnight at 4 °C with end-over-end rotation. Subsequently, the samples were centrifuged at 10,000 × *g* for 10 minutes at 4 °C. The supernatant containing the soluble protein fraction was collected, and the pellet discarded. Protein extracts were aliquoted and stored at - 80 °C until further use. Total protein concentration was determined spectrophotometrically on a DeNovix DS-11 Fx+ spectrophotometer (Wilmington, Delaware, USA) using the Pierce 660 nm assay (Thermo Fisher Scientific, Bremen, Germany).

### Proteomic analysis of a bark beetle protein extract by nano LC-MS/MS

Sample preparation for mass spectrometry was performed using a modified S-Trap micro spin column protocol (Protifi, Fairport, NY, USA). Briefly, 10 µg protein was reduced and alkylated with 200 mM tris(2-carboxyethyl)phosphine (TCEP) and 800 mM chloroacetamide (CAA) in 100 mM triethylammonium bicarbonate (TEAB) at a final ratio of 1:20 (v/v) in a total volume of 40 µl. After incubation for 30 min at 37 °C in the dark, SDS was added to a final concentration of 2%. Samples were then acidified with 12% aqueous phosphoric acid to a final concentration of 1%, mixed with S-Trap buffer (90% methanol, 100 mM TEAB; 6× sample volume), and loaded onto the column by centrifugation at 1000 × g for 1 min. Columns were washed six times with 150 µl S-Trap buffer, dried by centrifugation at 4000 × g for 1 min, and digested overnight at 37 °C with 1 µg Trypsin/Lys-C in 20 µl 50 mM TEAB. Peptides were sequentially eluted with 50 mM TEAB, 0.2% formic acid (FA), and 50% acetonitrile (ACN), dried, resuspended in 0.1% trifluoroacetic acid (TFA), and desalted using Pierce C18 spin tips (Thermo Fisher Scientific, Bremen, Germany) according to the manufacturer’s instructions. Desalted peptides were dried again, resuspended in 100 µl 0.1% TFA, and 3 µl were injected into a Q Exactive HF Orbitrap mass spectrometer (Thermo Fisher Scientific, Bremen, Germany) coupled to an Ultimate 3000 RSLC nano-HPLC system (Dionex). MS data acquisition and qualitative analysis using Proteome Discoverer v3.1.1.93 (Thermo Fisher, Vienna, Austria) were performed as described by Mayr et al., (2024). The resulting MS/MS spectra were searched against a curated *I. typographus* proteome database containing 23,923 entries (accessed January 2026) (Powell et al., 2021). Because bark beetle attacks are naturally associated with ophiostomatoid fungi that contribute to host colonization and modulation of tree defenses, searches were additionally performed against the Ophiostomataceae proteome database (27,636 entries) and the Ceratocystidaceae proteome database (15,273 entries) to assess the presence of potential fungal proteins in the beetle extracts. However, the present study was designed primarily to investigate beetle-derived elicitors under controlled conditions. The fungal databases were accessed through UniProt (https://www.uniprot.org/taxonomy/5152 and https://www.uniprot.org/taxonomy/1028423, respectively) in October 2025. Search parameters applied were as follows: enzyme trypsin (full); maximally 2 missed cleavage sites; 10Cppm precursor mass tolerance and 0.02CDa fragment mass tolerance; dynamic modifications allowed were oxidation/+15.995CDa (M), deamidation/+0.984CDa (N, Q), Gln->pyro-Glu/−17.027CDa (Q), and static modification carbamidomethylation/+57.021CDa (C). Proteins with more than two identified tryptic peptides, at least one of them unique, were reported.

### Plant material and bark beetle protein inoculation

Norway spruce seedlings (Provenance: Fi 89 (2.1/ts), 47.38°N, 12.3°E, 1450 m) were grown in a phytotron (Model D1200PLH, Aralab, Lisbon, Portugal) under controlled conditions (21 °C, 16 h light : 8 h dark photoperiod, 65% relative humidity, and approximately 150 µmol m_⁻_² s_⁻_ ¹ light intensity corresponding to ∼40% illumination). Seedlings were 10 weeks old at the time of treatment and approximately 8–10 cm in height. All sampling procedures were performed during the dark period, as the light cycle started at approximately 15:30 each day.

Three experimental conditions were established: (i) an untreated negative control, (ii) a mock challenge with buffer only (CHAPS or NP-40), and (iii) a challenge group treated with beetle protein extracts dissolved in the respective buffer (Figure 1B). Each treatment was applied along the stem surface using two different methods. In the first, buffer or protein solution was applied directly using a cotton swab (Q-tip). In the second, the same solutions were mixed with carborundum powder (10 mg mL⁻ ¹; silicon carbide powder, F320, ∼30 µm particle size, Werth-Metall, Germany) and gently rubbed mainly onto the central stem region using a Q-tip, resulting in mild abrasion of the outer stem surface and enhancing tissue contact. (Figure 1C). All seedlings remained under identical phytotron conditions throughout the experiment.

**Figure 1.**
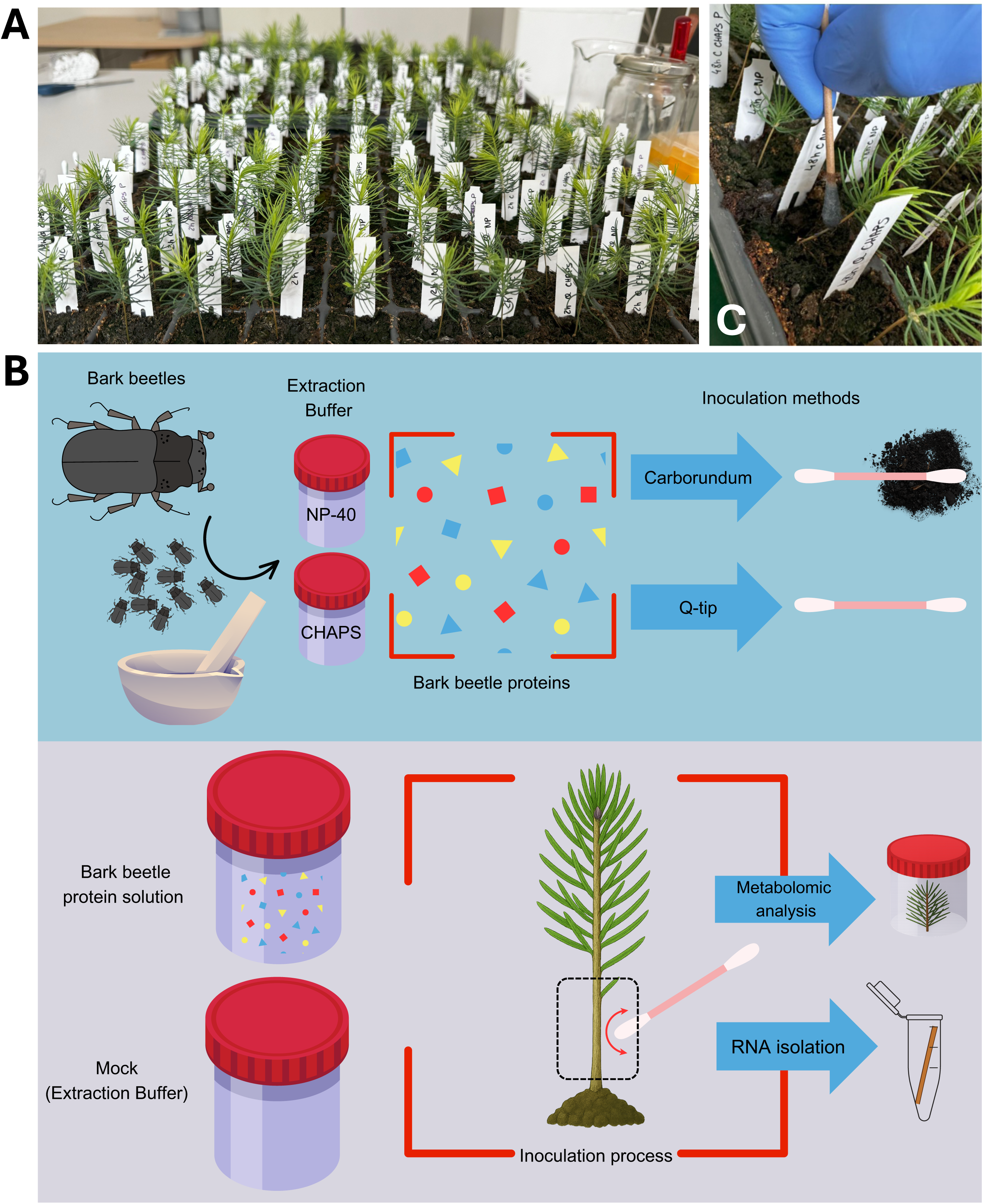
Phytotron-based setup to study the induced defense molecular reaction of Norway spruce to bark beetle (*Ips typographus*) protein extracts. **(A)** 10-week-old Norway spruce seedlings used for the experiment. **(B)** Schematic view of the experimental design, methodology used and sampling. **(C)** Bark beetle protein extract inoculation using Q-tip with carborundum.

The stems were harvested at two time points: 2Cand 48 hours post-inoculation (2 hpi; 48 hpi). At each time point, entire stem segments were collected from the region between the base of the stem and the start of the canopy and used for RNA extraction and transcriptomic analysis (Figure 1B). Four biological replicates were prepared per treatment and time point, with each replicate consisting of pooled stem tissue from two individual seedlings. In parallel, canopy tissue (consisting exclusively of needles, with no stem material) was harvested at both time points, and stored separately for targeted analyses of secondary metabolites. All samples were snap-frozen in liquid nitrogen and stored at −80C°C until further processing.

### RNA extraction and quality control

Frozen stems were cut into 1-cm segments and placed into 2-ml screw cap micro tubes (Sarstedt, Nümbrecht, Germany), prefilled with four 2.38-mm metal beads (Qiagen, Hilden, Germany) and homogenized using a MagNA Lyser (Roche, Rotkreuz, Switzerland). The tissue was disrupted in three cycles at 4,500 rpm for 15 s, with snap-freezing in liquid nitrogen between cycles. Care was taken to keep samples frozen before the addition of lysis buffer, as mechanical disruption in the absence of lysis buffer can result in RNA degradation due to released RNases.

Total RNA was extracted using the Spectrum™ Plant Total RNA Kit (Sigma-Aldrich, St. Louis, MO, USA), following Protocol B, with a few minor modifications. Briefly, Lysis Solution was freshly prepared by supplementing with 1% (v/v) 2-mercaptoethanol immediately prior to use. After homogenization, 500 µL of the supplemented Lysis Solution was added to each sample, followed by vortexing for 30 s and incubation at 56 °C for 5 min.

The clarified lysate was passed through a filtration column to remove debris, followed by binding of RNA to a silica column using the provided Binding Solution. After RNA binding, columns were washed and subjected to on-column DNase I digestion using the DNase I Digestion Set (Sigma-Aldrich) to eliminate residual genomic DNA. Columns were then washed three times with Wash Buffer II (Sigma-Aldrich, Vienna, Austria), instead of the standard two, to improve removal of potential contaminants. RNA was eluted in 50 µL of RNase-free water and stored at −80 °C until further processing. RNA concentration was measured using a DeNovix DS-11 FX+ spectrophotometer (DeNovix Inc, Wilmington, US). RNA integrity and quality were assessed using the Agilent RNA ScreenTape assay on a 4200 TapeStation system (Agilent Technologies, Vienna, Austria).

### Evaluation of differential gene expression through RT-qPCR

Twenty-four target genes were selected for RT-qPCR based on previous findings (Ramires et al. 2025). Elongation factor 1-α and polyubiquitin were included as reference genes as described in Ramires et al. (2025). Differentially expressed genes (DEGs) identified during Days 2 and 3 following a pioneer bark beetle attack in mature Norway spruce trees (n = 1,625) were retrieved from Ramires et al. 2025. From this set, DEGs with an adjusted p-value < 0.001 and a log2 fold change > 2 were selected (n = 208). We then focused on Day 2 and selected 18 genes with the lowest adjusted p-values for which primer design was feasible. In addition, six genes previously analyzed in the earlier trial (Ramires et al., 2025) were included using the same primer sets, as they have already been established as relevant markers of tree stress responses. Gene expression was normalized to untreated negative controls (NC), and statistical significance was determined by comparing each protein treatment with its corresponding buffer control. Primer sequences are listed in Supplementary Table S1 (Trujillo-Moya et al., 2020, 2022; Yakovlev et al., 2006, 2014, Ramires et al., 2025). The iScript Explore One-Step RT and PreAmp Kit (Bio-Rad, Hercules, CA, USA) was used for gDNA clearance, reverse transcription, and cDNA preamplification using 50 ng RNA input with 11 preamplification cycles, according to the recommended protocol for self-designed assay primers. The pre-amplified cDNA was diluted 1:10 in nuclease-free H_2_O for RT-qPCR. The 20 µL-PCR reactions included 1x HOT FIREPol EvaGreen qPCR Mix Plus ROX (Solis BioDyne, Tartu, Estonia), 200 nM of each primer and 2 μL diluted cDNA. All samples were analyzed in duplicates on an AriaMx Real-Time PCR system (Agilent) using the following temperature profile: 95 °C for 12 min, 40 cycles of 95 °C for 15 s and 60 °C for 1 min, followed by a melting curve step (60 °C–95 °C). Raw Cq-values were adjusted for PCR efficiencies and normalized to the geometric mean of both reference genes. Expression changes relative to the negative control groups were calculated with the comparative ddCT method (Livak and Schmittgen, 2001). Statistical analyses (unpaired t-test with Welch’s correction) were calculated using GraphPad Prism 10.1.3 (GraphPad Software, San Diego, CA, USA).

### RNA-seq differential expression analysis

Extracted RNA concentration and quality were assessed with NanoDrop 2000C (Thermo Fisher Scientific, Vienna, Austria) and Fragment Analyzer (Agilent, Vienna, Austria) for NC 2 h / 48 h, Mock 2 h / 48 h and Protein 2 h / 48 h samples. For those, indexed libraries were prepared from 100 ng of RNA using the QuantSeq 3’ mRNA-seq Library Prep Kit REV (Lexogen, Vienna, Austria) for Illumina (015UG009V0241) following the manufacturer’s instructions and analysed with Fragment Analyzer (Agilent) using the HS-DNA assay. Libraries were quantified using a Qubit dsDNA HS assay kit (Thermo Fisher Scientific, Waltham, MA, USA) and sequenced on Illumina NovaSeq6000 with a SR100 read mode (Illumina, San Diego, CA, USA). Sequencing quality control of the raw reads was assessed using FastqQC software and adapter sequences were removed with cutadapt (v1.18; (Martin, 2011)). Alignment to the reference genome of *P. abies* (Nystedt et al., 2013) and read counting were performed using STAR (v2.6.1a; (Dobin & Gingeras, 2015)) and featureCounts (v1.6.4; (Liao et al., 2014)), respectively. RSeQC (Wang et al., 2012, 2016) was used to assess the quality of data after alignment. DEGs were identified with DESeq2 (v1.18.1; (Love et al., 2014)), using Mock samples as controls and adjusting p-values for multiple comparisons with the FDR method. Genes were identified as differentially expressed if they exhibited a minimum 2-fold change and had adjusted p-values below 0.05.

### GO enrichment and KEGG pathways analysis

At each time point, DEGs were subjected to gene ontology (GO) enrichment analysis. GO annotations were retrieved from PlantGenIE (https://plantgenie.org/), and enrichment was calculated using the topGO package in R (v2.48.0; Alexa & Rahnenfuhrer, 2023) with the *weight01* algorithm and a one-sided Fisher’s exact test. GO terms with p-values below 0.01 were considered significantly enriched. Results were visualized using ggplot2 (v3.4.2; Wickham, 2009).

For Kyoto Encyclopedia of Genes and Genomes (KEGG) pathway analysis, we applied the same workflow as previously described in Trujillo-Moya et al., 2020 and 2022. DEGs were first assigned to KEGG orthologs using a bidirectional best hit (BBH) search against a database of 67 plant species (Supplementary Table S2). The number of genes mapped to each pathway was then summarized with custom Python scripts (Ahmad et al., 2025), separately for up- and downregulated genes at each time point.

### Comparison with field experiment

DEGs obtained here were compared to our previous molecular study on the modulated defense response after pioneer bark beetle attack on 35-year-old clonal trees on a controlled infestation field experiment (Ramires et al., 2025). To assess whether DEG overlap between controlled-environment (Phytotron) and field (Forest) conditions exceeded random expectations, Fisher’s Exact Test was applied across all pairwise comparisons between Phytotron samples collected at 2 and 48 h post-inoculation and Forest samples collected at 2, 3, 5, and 7 days following pioneer bark beetle attack. Up- and down-regulated DEGs were analyzed separately to preserve biological directionality, yielding comparisons of up-regulated genes against up-regulated genes and down-regulated genes against down-regulated genes across all time point combinations. This resulted in eight pairwise comparisons in total (two Phytotron time points × two Forest time points × two directions). For each comparison, a 2 × 2 contingency table was constructed as follows: (i) genes present in both the Phytotron and Forest sets (overlap), (ii) genes present only in the Phytotron set, (iii) genes present only in the Forest set, and (iv) genes absent from both sets. The total gene universe used as background comprised 37,565 genes in the reference genome. Fisher’s Exact Test was applied one-sided (alternative hypothesis: the observed overlap is greater than expected under independence) to test for significant enrichment of shared DEGs. The odds ratio was used as a measure of effect size. To account for multiple comparisons across all eight tests, p-values were adjusted using the Benjamini–Hochberg procedure (Benjamini & Hochberg, 1995) controlling the false discovery rate (FDR). An adjusted p-value threshold of 0.05 was used to determine statistical significance. All analyses were performed in R (version 4.5.2) (R Core Team, 2022).

### Targeted analyses of secondary metabolites

Due to the limited amount of stem tissue available for RNA isolation, needle samples were used as a practical proxy to assess changes in secondary metabolites and to test for systemic defense response. Whereas transcriptomic analyses were performed on treated stem tissue, targeted metabolomic analyses were conducted on needles to investigate distal metabolic responses following stem exposure to bark beetle-derived proteins.

Sample preparation, extraction and analysis of phenolic metabolites were conducted as previously described (Ganthaler et al., 2017). Briefly, freeze-dried samples were homogenized with a ball mill (Tissuelyzer II, Qiagen, Hilden, Germany) for 2 min at 30 Hz and phenolics were extracted from 12.0 ± 0.2 mg of powder in two successive steps each using 1 ml of 95% (v/v) ethanol, containing orientin, naringin, and pinosylvin as internal standards. Twelve flavonoids (apigenin, eriodictyol, kaempferol, kaempferol 3-glucoside, kaempferol 7-glucoside, kaempferol 3-rutinoside, quercetin, quercetin 3-glucoside, naringenin, taxifolin, catechin, gallocatechin), four stilbenes (astringin, isorhapontin, piceid, piceatannol), two simple phenylpropanoids (picein, gallic acid), a diterpenoid (abietic acid) and a precursor (shikimic acid), were identified and quantified by liquid chromatography-mass spectrometry (UHPLC-MS). Analytes were separated on a reversed-phase column (NUCLEODUR C18 Pyramid, EC 50/2, 50C×C2 mm, 1.8 μm, Macherey–Nagel, Düren, Germany) using an Eksigent ultraLC 100 UHPLC system and detected with a QTRAP 4500 mass spectrometer (AB SCIEX, Framingham, MA, USA) operated in negative ion mode using scheduled multiple reaction monitoring (sMRM) with the Analyst software (version 1.71, AB SCIEX). Commercial standards were used to establish calibration curves for absolute quantification. Peaks integration, normalization relative to internal standards and concentration calculations were performed using the SCIEX OS-MQ software (version 1.6.1, AB SCIEX).

## Results

### Bark beetle protein extracts consist primarily of beetle-derived proteins

In total, 1,582 unique proteins were identified from the bark beetle extracts across both extraction conditions. Of these, 1,559 proteins were detected in the CHAPS extract and 1,527 in the NP-40 extract, indicating slightly broader proteome coverage with CHAPS. A total of 1,495 identical protein entries were shared between both extraction conditions, corresponding to 95.9% of the CHAPS proteome and 97.9% of the NP-40 proteome, while 64 proteins were unique to CHAPS and 32 to NP-40. Only a minor fraction of the identified proteins was of fungal origin (<0.2%). In each extract, six proteins were attributed to Ophiostomataceae and one to Ceratocystidaceae. The Ceratocystidaceae-derived protein was detected with both extraction methods, whereas within the Ophiostomataceae-assigned proteins, two were specific to CHAPS and two were specific to NP-40, leaving four shared between both extracts. Overall, both detergents yielded highly comparable proteomes composed mostly of bark beetle-derived proteins, with only negligible fungal contribution (Supplementary Table S3).

### RT-qPCR showed that NP-40 extract inoculation effectively induces a defense response in Norway spruce seedlings

RT-qPCR analysis of 24 selected genes validated expression patterns following bark beetle protein extract inoculation in Norway spruce seedlings. Responses were gene-and condition-specific, with significant regulation occurring more frequently at 2 h than at 48 h, indicating an early and transient response. Both CHAPS- and NP-40-based treatments induced 17 genes, although NP-40 treatments at 2 h showed more consistent upregulation in the presence of protein extract, particularly when applied with carborundum (Supplementary Figure S1). At 2 h, genes related to plant-pathogen interaction and MAPK signaling, including Calmodulin-like 43 (MA_18054g0010), MPK6 (MA_10437020g0010), and MYC2 (MA_10435905g0020), were significantly upregulated compared with buffer controls (Figure 2A). At 48 h, additional genes linked to MAPK signaling (Basic Endochitinase, MA_8921185g0010), hormone signaling (JAZ, MA_9455802g0010), and secondary metabolism, including phenylpropanoid (pDIR2, MA_722981g0010), flavonoid (Chalcone and stilbene synthase family protein, MA_57359g0010), and terpenoid biosynthesis (Terpene synthase, MA_70145g0010), were mainly induced.

**Figure 2.**
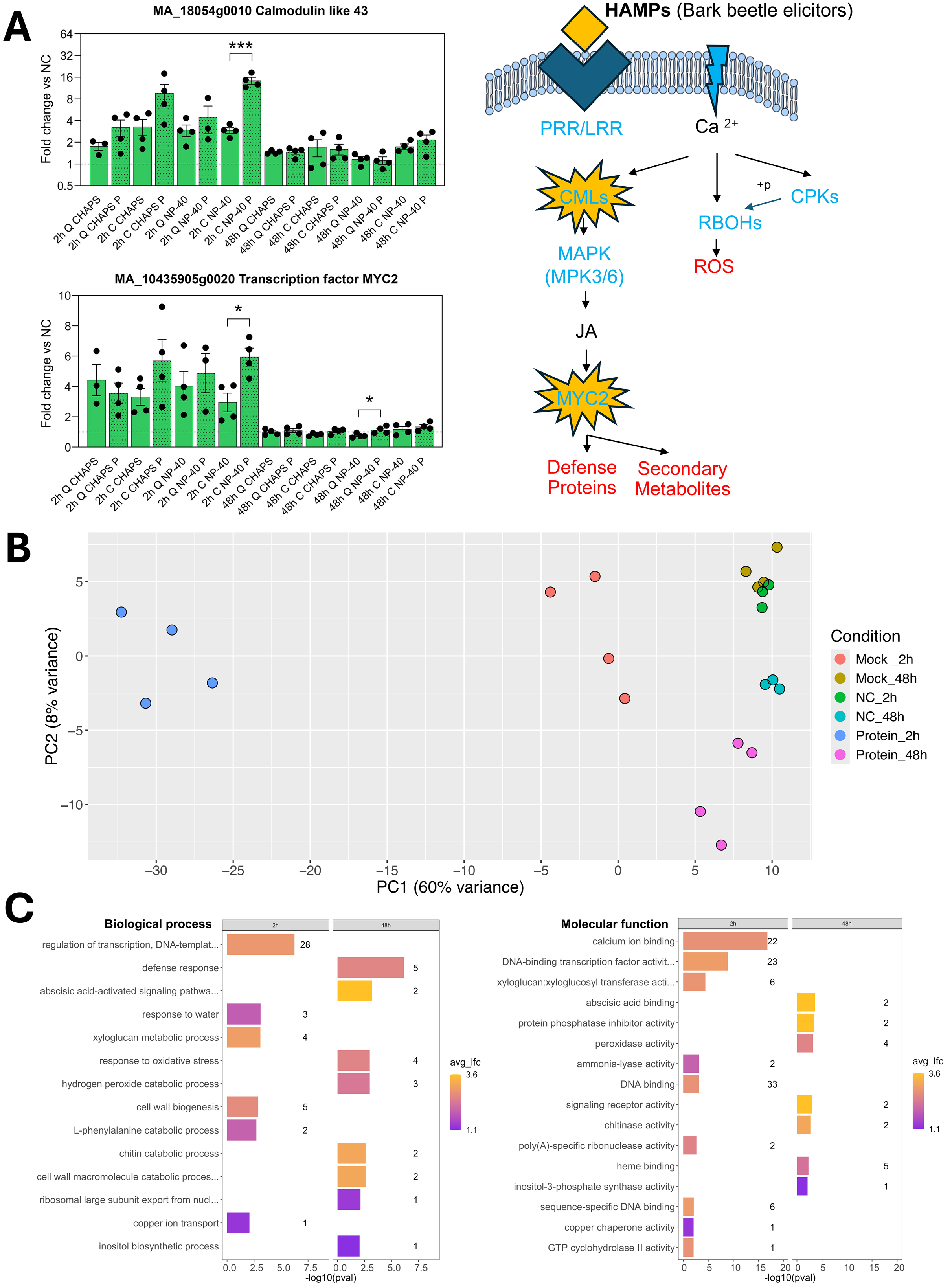
Induced molecular defense responses in Norway spruce triggered by bark beetle (*Ips typographus*) protein extracts. **(A)** RT-qPCR–monitored expression of a calcium-binding protein CML (plant–pathogen interaction pathway) and the transcription factor MYC2 (MAPK and hormone signaling pathways) with a schematic representation of a putative HAMP-triggered immunity-like response **(B)** PCA of gene expression profiles from RNA-seq when NP-40 protein extract was applied by using Q-tip plus carborundum. **(C)** GO enrichment on biological processes and molecular functions for the over-expressed DEGs sets both at 2 h and 48 h.

Under NP-40 protein extract treatments, ABC transporter (MA_6870g0010), Chalcone and stilbene synthase family protein (MA_10426246g0020), and Phenylalanine ammonia-lyase 4 (MA_44561g0010) were upregulated, whereas the latter two, together with Flavanone 3-hydroxylase (MA_10435304g0010), were downregulated when using CHAPS protein extracts. Conversely, four genes were uniquely induced by CHAPS protein extract treatments, including Jasmonate-zim-domain protein 12 (MA_10433497g0010) and Chalcone–flavanone isomerase (MA_50294990g0010). Several genes showed no significant differential expression under any treatment or time point, suggesting limited involvement in the early defense response.

### RNA-seq reveals biphasic immune responses and conserved defense signatures

Induced defense response triggered by NP-40 protein extract was further explored and a principal component analysis (PCA) of the RNA-seq differential expression data (Figure 2B; Supplementary Table S4) revealed clear separation between the Protein and Mock/NC groups. PC1 and PC2 together explained 68% of the total variance, with PC1 accounting for 60%. In total, 488 and 84 genes were differentially expressed between Protein and Mock at 2 and 48 hpi, respectively, of which on average 79% were over-expressed and 21% were under-expressed. Enriched GO terms were identified primarily for differentially over-expressed genes at each time point, with no shared terms observed between time points (Figure 2C). At 2 hpi, the enriched terms were associated with transcriptional regulation (e.g., DNA-binding transcription factor activity, DNA binding), xyloglucan metabolic process, cell wall biogenesis, L-phenylalanine catabolism (ammonia-lyase activity), and calcium signaling (calcium ion binding). In contrast, at 48 hpi, enriched terms were related to defense responses (e.g., defense protein activity), chitin catabolic processes (chitinase activity), abscisic acid-activated signaling pathways, and responses to oxidative stress (peroxidase activity). The limited overlap between DEGs at 2 and 48 hpi further supported a temporally structured defense response (Figure 3). Nevertheless, comparative analysis with our previous field experiment at 2 and 3 dpi showed that, on average, approximately 30% of the DEGs identified here overlapped with those detected in field-grown Norway spruce trees exposed to *I. typographus* attack (Ramires et al., 2025). This overlap was significant and unlikely to have occurred by chance (Supplementary Table S5). Among upregulated DEGs, 29% and 53% were shared at 2 h and 48 h, respectively, whereas 21% and 25% of downregulated DEGs overlapped at the corresponding time points (Figure 3).

**Figure 3.**
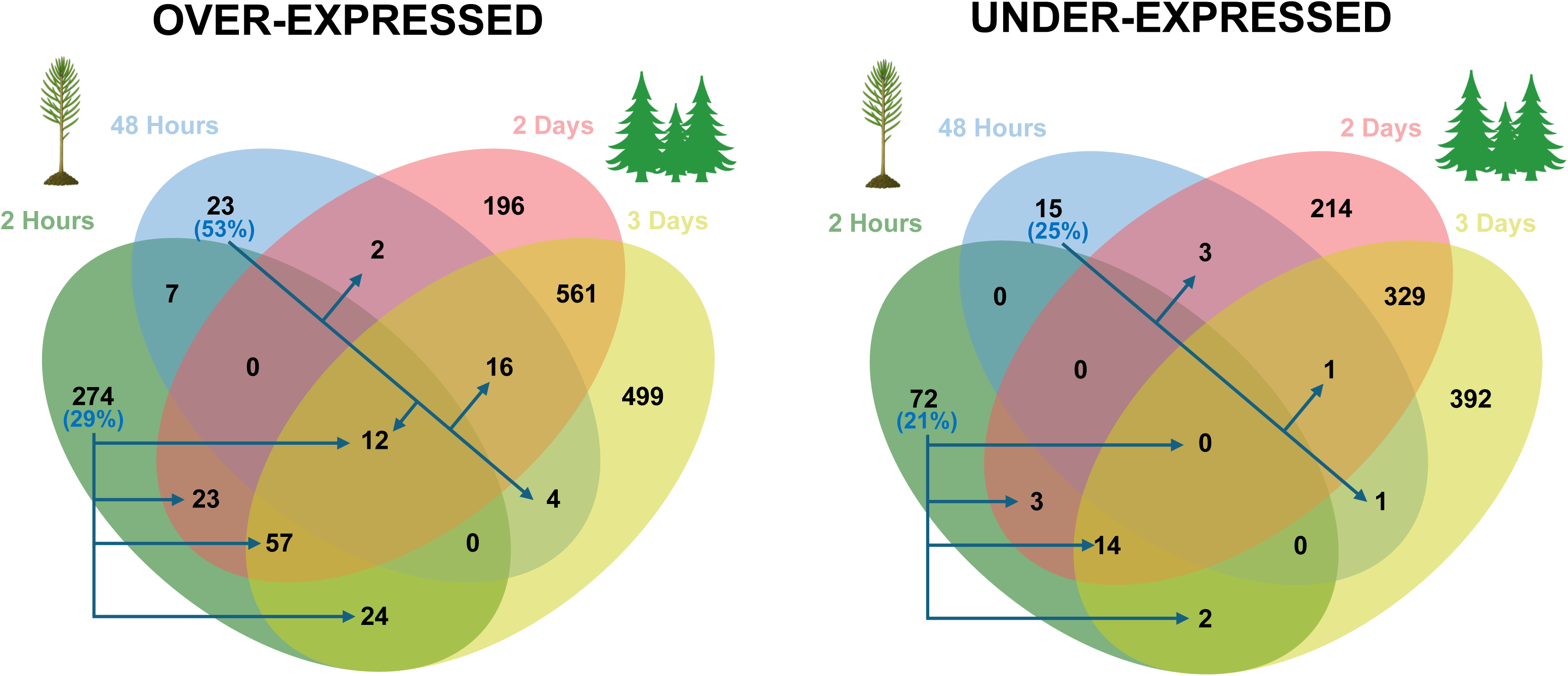
Comparison between DEGs in this 10-week-old seedling phytotron setup (2 and 48 hours) vs field setup (2 and 3 days; Ramires et al., 2025). Venn diagrams for over and under-expressed DEGs shared by both approaches including the percentage overlap.

KEGG annotation, covering 33% of DEGs, revealed that at 2 h most upregulated genes belong to pathways related to Norway spruce defense, particularly signal transduction and environmental adaptation, including plant–pathogen interaction, MAPK signaling, and plant hormone signal transduction (Table 1). This early response was characterized by the induction of key signaling components such as calcium-binding proteins (e.g., CMLs), a calcium-dependent protein kinase (CPK), and a respiratory burst oxidase (RBOH), followed by MAPK-associated regulators (e.g., MYC2) and jasmonate signaling elements (e.g., JAZ). In parallel, genes involved in secondary metabolism (e.g., PAL) and in jasmonic acid (JA) biosynthesis via the octadecanoid pathway from α-linolenic acid were also induced. At 48 h, KEGG annotation identified additional defense-related genes, including well-known defense proteins such as basic endochitinase B, peroxidases associated with the phenylpropanoid biosynthetic pathway, and enzymes of the terpenoid backbone biosynthesis pathway, such as phosphomevalonate kinase and isoprene synthase, both involved in monoterpene biosynthesis. Most of these KEGG-annotated genes detected at both 2 h and 48 h were also shared with adult trees and complemented the schematic representation of a putative HAMPs-triggered immunity-like response (Figure 2A; Supplementary Figure S2G)(Ramires et al., 2025).

**Table 1.** List of Differentially Expressed Genes (DEG) belonging to the main defense related KEGG pathways.

Additional genes not assigned to KEGG pathways, representing 67% of DEGs, included multiple receptor classes involved in immune perception, dominated by leucine-rich repeat receptor-like kinases (LRR-RLKs) and intracellular NLR proteins, suggesting the activation of both surface-mediated perception of herbivore-associated molecular patterns (HAMPs) and intracellular immune signaling pathways (Supplementary Table S4). This was accompanied by the overexpression of numerous transcriptional regulators, including MYB, WRKY, and Ethylene Responsive Factor (ERF) transcription factors, among others. Early induction of defense-related proteins was also evident, including chitinase-2-like, alongside genes associated with phenolic metabolism and structural reinforcement such as scopoletin glucosyltransferase and dirigent proteins among others. At 48 h, the transcriptional profile shifted toward the accumulation of defense proteins, including Defensin-like proteins, PR10 proteins, thaumatin (PR-5) and a proteinase inhibitor potentially targeting digestive proteases in the bark beetle gut, together with enzymes linked to cell wall modification and secondary metabolism, such as laccase 12, additional dirigent proteins, and a sesquiterpene synthase (zerumbone synthase). Most of these genes were also differentially expressed in older trees following bark beetle attack (Ramires et al., 2025), suggesting that key immune and metabolic defense components are conserved between seedlings and mature trees. (Supplementary Table S4). Overall, the transcriptomic response shifted from early immune signaling and transcriptional reprogramming at 2 h to defense protein accumulation and structural and chemical reinforcement at 48 h.

### Profiling of phenolics suggests limited systemic defense responses in needles

Because stem tissue was largely consumed for transcriptomic analyses, profiling of 23 phenolic compounds was performed on the remaining needle tissue to assess whether transcriptional changes in the stem were associated with systemic metabolic responses in needles. For most compounds measured, no differences were observed between seedlings treated with the protein extract and those with the respective buffer, suggesting that the systemic needle response was limited within the 48-hour window (Supplementary Figure S3). However, a subset of metabolites exhibited treatment- and time-specific accumulation changes. At the 2-hour time point, no consistent or significant differences were observed between protein-treated and buffer-treated samples for any of the measured compounds. At 48 hours, several compounds showed visible trends, though most changes were not statistically significant due to high variation across biological replicates. Flavonoid aglycones (quercetin, naringenin, kaempferol) and gallic acid tended to exhibit lower levels in protein-treated samples compared with buffer-only or control groups. In contrast, kaempferol glycosylated forms (kaempferol-3-glucoside, kaempferol-7-glucopyranoside, and kaempferol-3-rutinoside) displayed a tendency toward higher accumulation in the protein-treated group at 48 hours. All other compounds remained unchanged across treatments and time points.

## Discussion

Our proof-of-concept approach showed that protein extracts derived from *I. typographus* adults can effectively trigger measurable defense responses in *P. abies* seedlings under controlled laboratory conditions. This assay bypasses the need for live insect infestation and provides a scalable and reproducible platform to study early molecular responses, in line with the need for more standardized insect-derived elicitor systems (Erbilgin et al., 2006; Zhao et al., 2010).

The diversity of proteins recovered from *I. typographus* adult beetles seems to be only marginally influenced by the extraction buffer used. While with CHAPS a slightly higher number of identified proteins could be observed than with NP-40, both extracts showed a substantial overlap, with 1495 identical protein entries shared between conditions. Nevertheless, the minimal increase in proteome coverage with CHAPS did not correspond to stronger biological activity in the downstream seedling assays. On the contrary, NP-40 extracts elicited a more pronounced transcriptional defense response in Norway spruce, suggesting that elicitor activity depends less on the total number of recovered proteins than on the extraction or preservation of specific bioactive components. This points to qualitative differences between the buffer-specific fractions, with NP-40 likely recovering a protein subset more relevant for host recognition and defense activation or increasing the tree’s tissue permeability for this specific protein cargo. Although fungal associates of *Ips typographus* are important contributors to host colonization and tree defense responses during natural attacks, fungal proteins represented only a negligible proportion of the identified proteins in both extracts. The differential biological activity observed here is therefore most plausibly explained by differences in bark beetle-derived protein composition. Nevertheless, the specific elicitor(s) responsible for triggering these responses remain unresolved.

The transcriptional patterns observed in the RT-qPCR analysis align with those previously reported under biotic stress in RNA-seq studies (Trujillo-Moya et al., 2020, 2022, Ramires et al., 2025), supporting the reproducibility and reliability of the selected gene pool under controlled elicitor treatments in Norway spruce seedlings, especially when NP-40 was used in combination with carborundum as an abrasive to facilitate protein extract penetration into stem tissue. In the carborundum treatment, the protein extract was gently rubbed across the stem surface using a cotton swab.

This resulted in mild abrasion and partial removal of superficial stem surface layers, likely enhancing contact between the applied protein extract and underlying outer stem tissues.

RNA-seq results on NP-40 elicited samples confirmed what was glimpsed by RT-qPCR, that the exposure of plants to the insect protein extract triggered a temporally structured immune response characterized by an early signaling phase (2 h) followed by a later execution phase (48 h), as clearly evidenced by the GO enrichment analysis, which revealed no shared terms between the two time points.

At 2 h, the transcriptional profile was dominated by genes associated with perception, early signal transduction, and transcription, suggesting rapid activation of defense signaling pathways. In addition, strong induction of calcium-binding proteins, calcium-dependent kinases, MAPKKKs, and respiratory burst oxidase homologs (RBOHs) was observed, indicating activation of calcium influx and ROS-mediated signaling. This early signature is consistent with a PTI-like response, in which recognition of herbivore-associated molecular patterns (HAMPs) or damage-associated signals initiates rapid defense signaling cascades. The concurrent upregulation of key transcription factors such as WRKYs, ERFs, and MYC2, together with activation of jasmonate (AOS, LOX, OPR) and ethylene-related pathways, further supports the establishment of a JA/ET-dominated transcriptional reprogramming typically associated with herbivory. The induction of MAPK6 and JAZ domain proteins is consistent with the early involvement of jasmonate- and calcium-mediated signaling cascades in conifer defense against herbivores (Franceschi et al., 2005; Hammerbacher et al., 2019; Wilkinson et al., 2022). Although several NLR-related genes were induced, the use of a bark beetle protein extract does not allow us to infer canonical effector-triggered immunity (ETI), as no specific effector-receptor interaction can be resolved in this system. Instead, the observed transcriptional changes likely reflect broader defense activation in response to insect-derived elicitors. Even with carborundum-assisted inoculation, the use of protein extracts does not recapitulate the active and specific intracellular delivery of intact effectors required for ETI activation. In contrast, at 48 h the transcriptional landscape shifted toward the accumulation of defense proteins. These included pathogenesis-related proteins (PR10, thaumatin/PR-5), defensins, chitinases (Liu & Ekramoddoullah, 2006; Van Loon & Van Strien, 1999), a proteinase inhibitor that might inhibit digestive proteases in the bark beetle gut, as well as enzymes involved in ROS metabolism and cell wall reinforcement (peroxidases, laccases, dirigent proteins).

Upregulation of genes related to secondary metabolism, such as phenylpropanoid pathway enzymes (2 h) and terpene synthases (48 h), indicates the consolidation of chemical and structural defenses, as well as antimicrobial activity (Dao et al., 2011; Hammerbacher et al., 2019; Treutter, 2006). However, the flavonoid pathway was not significantly affected, contrasting with field observations reported by Ramires et al. (2025), where a marked over-expression was detected. The quantification of selected phenolic compounds did not reveal consistent compound-level changes, although substantial variability was observed across biological replicates, particularly at 48 h. This variability may reflect differences in systemic signaling responses among seedlings. For instance, the few phenylpropanoids and terpene profiles analyzed here did not reflect the induction patterns suggested by the DEGs. Notably, several flavonoid compounds (quercetin, naringenin, kaempferol, and gallic acid) tended to decrease in the presence of beetle protein extracts, while certain kaempferol glycosides showed slight increases in protein-treated samples. This overall reduction in metabolite abundance could indicate a reallocation of metabolic resources toward the local site of attack in the stem, where defense responses are most immediately required, potentially limiting the accumulation of secondary metabolites in distal tissues such as the canopy.

Given that localized defense induction often precedes systemic signaling, more time or additional stimuli may be required for a systemic response to manifest (Franceschi et al., 2005). Alternatively, protein-specific signaling could actively repress certain systemic responses as part of a more complex regulatory mechanism (Pieterse et al., 2012). Moreover, variable systemic responses are common, as observed in other plants responding to localized herbivory, and can be affected by individual variability (Arimura et al., 2005). Altogether, this suggests that metabolite analyses should be performed directly in stem tissue to properly capture the local metabolic changes induced by this biotic stress. Increasing the number of seedlings in the experimental design would ensure sufficient plant material for a robust analysis on this and other omics.

Collectively, these results support a model in which insect-derived proteins first activate a rapid perception and signaling phase (2 h), followed by a later execution phase (48 h) characterized by accumulation of transcripts coding for defense proteins. This temporal progression is consistent with an herbivory-induced defense program that shares core components with canonical plant immune responses to microbial pathogens. Differences between field and phytotron conditions must be interpreted cautiously, as they may arise from multiple interacting factors, including age, genotype, clone versus provenance effects, developmental stage, sampling time point, tissue specificity, inoculum load, and mechanical damage. These variables likely contribute substantially to the discrepancies observed between controlled and natural environments. In addition, the remarkable complexity of the Norway spruce genome, as highlighted by Nystedt and more recently by Cui, makes comprehensive transcriptomic interpretation challenging (Cui et al., 2021; Nystedt et al., 2013). Future studies incorporating spatial transcriptomics could provide cell-type-specific resolution of gene expression, enabling a more precise understanding of tissue-level responses during these interactions. While the use of clonal material will ensure experimental consistency (e.g., somatic embryogenic clones), future work should also consider variation among different beetle populations, as genetic diversity within *I. typographus* may influence elicitor composition and consequently the host defense response (e.g., Mykhailenko et al., 2024).

In summary, this study demonstrates that protein extracts from *I. typographus* can trigger biphasic immune responses in *P. abies* seedlings that recapitulate key aspects of bark beetle-induced defenses observed in mature trees. Although this system does not fully reproduce the ecological complexity of natural infestations, it provides a robust and reproducible platform for dissecting elicitor-driven conifer defense responses under controlled conditions. Because bark beetles threaten a wide range of forest tree species worldwide, this elicitor-based approach may also offer broader applicability for comparative studies of bark beetle-induced immunity across conifers.

**Supplementary Figure S1.** RT-qPCR on selected 24 defense related genes. Name of each condition is: 1) time point (2 h / 48 h), 2) inoculation method (Q: Q-tip alone / C: Q-tip plus Carborundum), 3) lysis buffer (CHAPS / NP-40), 4) when protein extract was used (P). Dotted line represents expression levels in untreated negative controls.

**Supplementary Figure S2.** KEGG pathways related to plant defense. **(A)** Plant-pathogen interaction. **(B)** MAPK signaling pathway-Plant. **(C)** Plant hormone signal transduction. **(D)** Alpha-Linolenic acid metabolism. **(E)** Phenylpropanoid biosynthesis. **(F)** Terpenoid backbone biosynthesis. **(G)** Schematic representation of a putative HAMP-triggered immunity-like response, now expanded with the RNA-seq results revealing a broader set of over-expressed DEGs.

**Supplementary Figure S3.** Comparative analysis of phenolic compounds in 10-week-old Norway spruce needles sampled 2 h and 48 h after protein inoculation. (A, B) Principal component analyses score plots generated from the relative abundances of 21 compounds detected in needles from the negative control (black), mock treatment (orange) and NP-40 protein extract treatment (red) at 2 h (A) and 48 h (B). (C) Hierarchically clustered heatmap illustrating the impact of the treatment on the accumulation of each detected phenolic compound at 2 h and 48 h. Data are log2 ratios of the mean metabolite relative abundances under each treatment relative to the corresponding negative controls, at the same time point (n = 4). Stars indicate Benjamini–Hochberg adjusted p-values < 0.05 and circles < 0.1, displayed in black for the comparison to the negative controls and in white for the comparison between the mock and protein extract treatment.

**Supplementary Table S1.** List of primers for RT-qPCR. Target gene information, primer sequences, and RT-qPCR assay details.

**Supplementary Table S2.** Sequence datasets of plant species applied for KEGG Orthology (KO) assignment using the KEGG Automatic Annotation Server (KAAS). KEGG Orthology (KO) assignment was applied using the Bi-directional Best Hit (BBH) method and all datasets of dicot plants, monocot plants and a basal magnoliophyta were selected as reference organisms.

**Supplementary Table S3.** Pseudo-quantitative proteomic profiling of *Ips typographus* protein extracts obtained using NP-40 and CHAPS extraction buffers

**Supplementary Table S4.** RNA-seq differentially expressed genes (DEGs). **(A)** DEGs summary for both 2 h and 48 h when log2FoldChange (≥ 1 or ≤ −1) and padj (≤ 0.05) filter criteria was applied. **(B)** 2 h DEGs Protein vs Mock. **(C)** 48 h DEGs Protein vs Mock**. (D)** KEGG pathway fingerprints for both DEGs sets, numbers represent the key enzymes from each pathway that are differentially expressed.

**Supplementary Table S5.** Overlap of differentially expressed genes (DEGs) between controlled (Phytotron) and field (Forest) conditions. DEGs from this study were compared with those from a previous field experiment (Ramires et al., 2025) across two time points in each condition, analyzing up- and down-regulated genes separately (eight pairwise comparisons). Overlap significance was tested using one-sided Fisher’s Exact Test (greater-than-expected overlap) based on 2 × 2 contingency tables and a background of 37,565 genes. The table shows gene set sizes (N), observed and expected overlaps, odds ratios, raw p-values, and Benjamini–Hochberg adjusted p-values (FDR). Significant enrichment was detected in most comparisons, indicating strong concordance between conditions. Significance codes: *** p < 0.001; ** p < 0.01; * p < 0.05; ns, not significant.

## Acknowledgements

This research was supported using resources of the VetCore Facility (Molecular Biology and Mass Spectrometry) of the University of Veterinary Medicine Vienna. Libraries were sent to the Lexogen NGS Services (Vienna, Austria) for sequencing.

## Author contributions

The project was conceptualized by CTM. The research was designed by MR and CTM. Bark beetle rearing and cage setup carried out by SN and MS. The experiment was conducted by MR. Sample processing was performed by MR. RNA isolation and RT-qPCR analysis were conducted by RE. GO enrichment analysis, KEGG pathway analysis and Fisher’s Exact Test were carried out by AM. Proteome extraction and LC-MS data analysis were performed by MR. Identification and quantification of phenolic compounds, diterpenes, and VOCs were carried out by EA. Data analysis and initial manuscript drafting were done by MR and CTM. Research supervision and project coordination was provided by CTM. CTM and MvL were responsible for funding acquisition. All authors edited and approved the manuscript.

## Data availability

Raw sequence data have been submitted to the NCBI Short Read Archive (SRA) under BioProject number PRJNA1470582.

The mass spectrometry proteomics data has been deposited to the ProteomeXchange Consortium via the PRIDE (Perez-Riverol et al., 2022) partner repository with the dataset identifier PXD078890 and 10.6019/PXD078890

