## Supplementary figures and images for "Bark beetle protein elicitors trigger biphasic immune responses in Norway spruce seedlings"

### Supplementary Figure S1

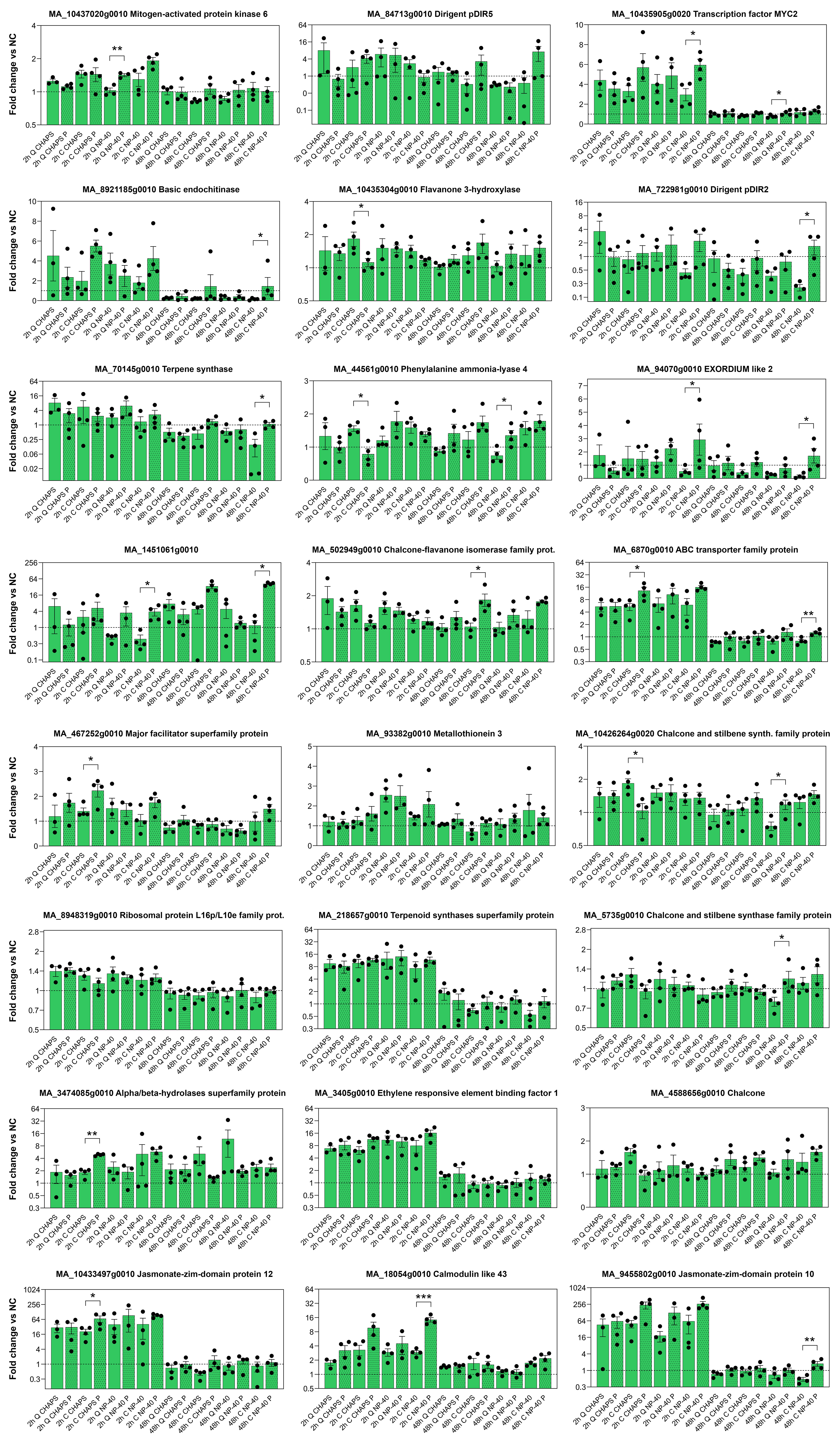

### Supplementary Figure S2

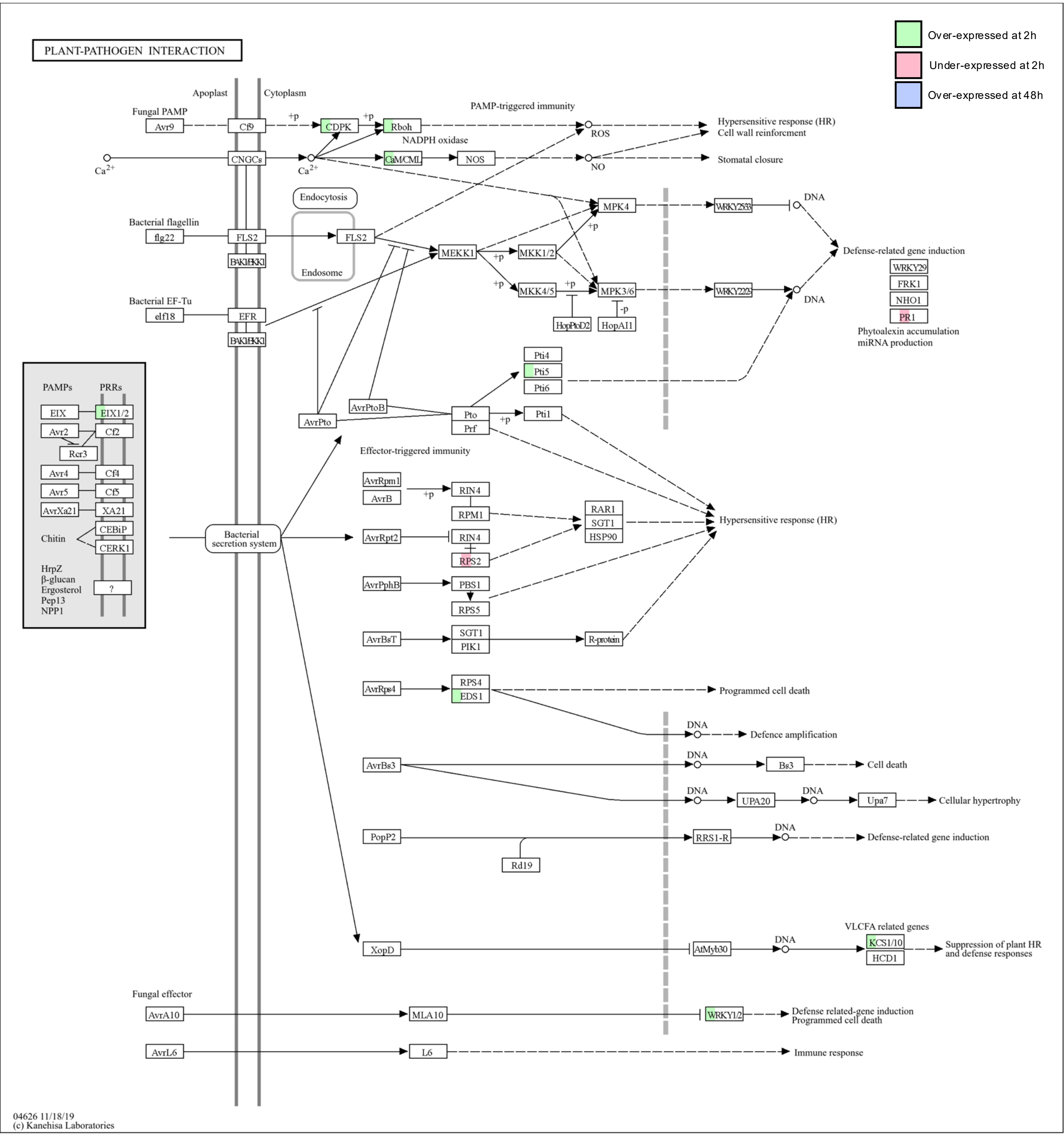

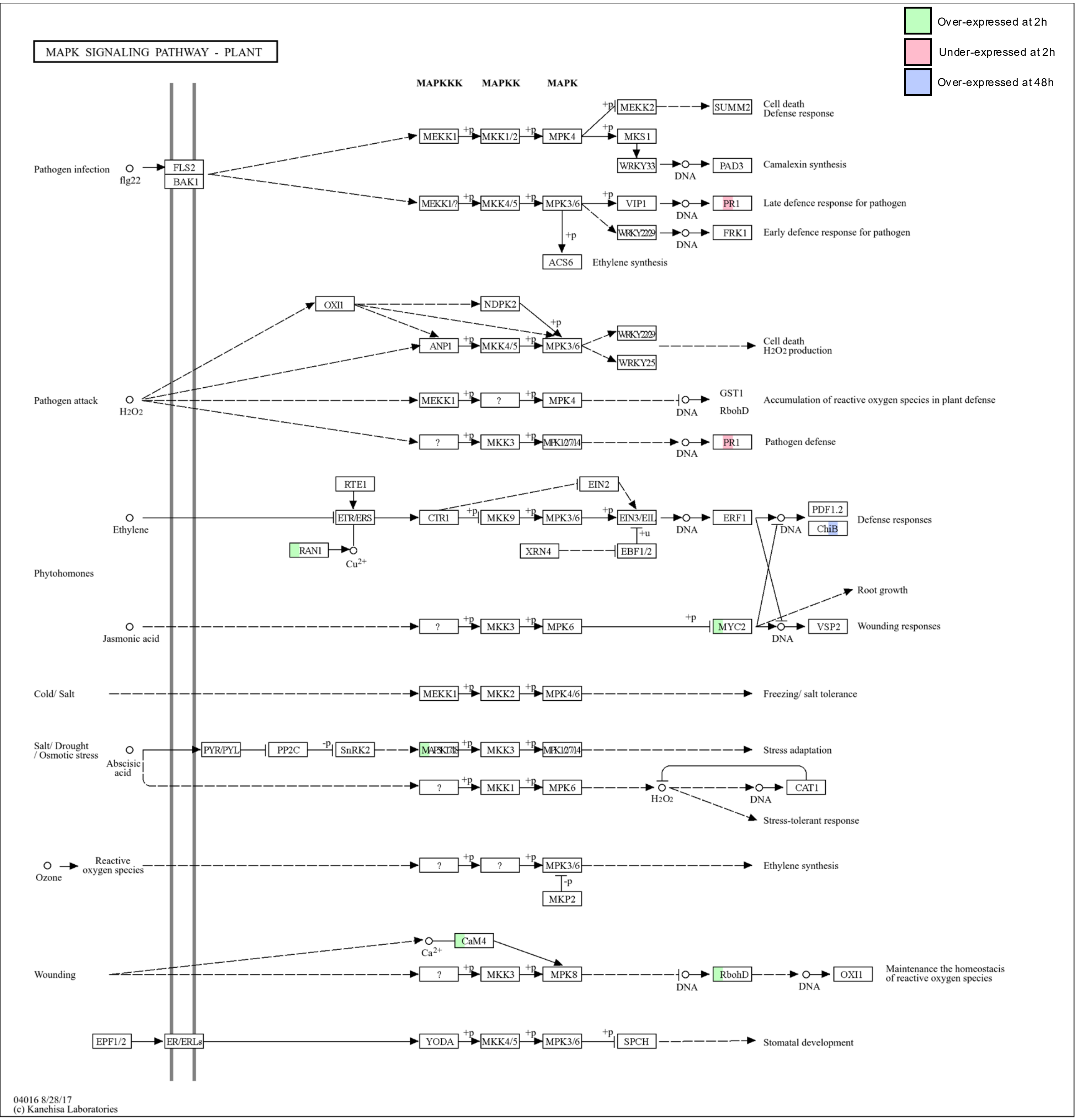

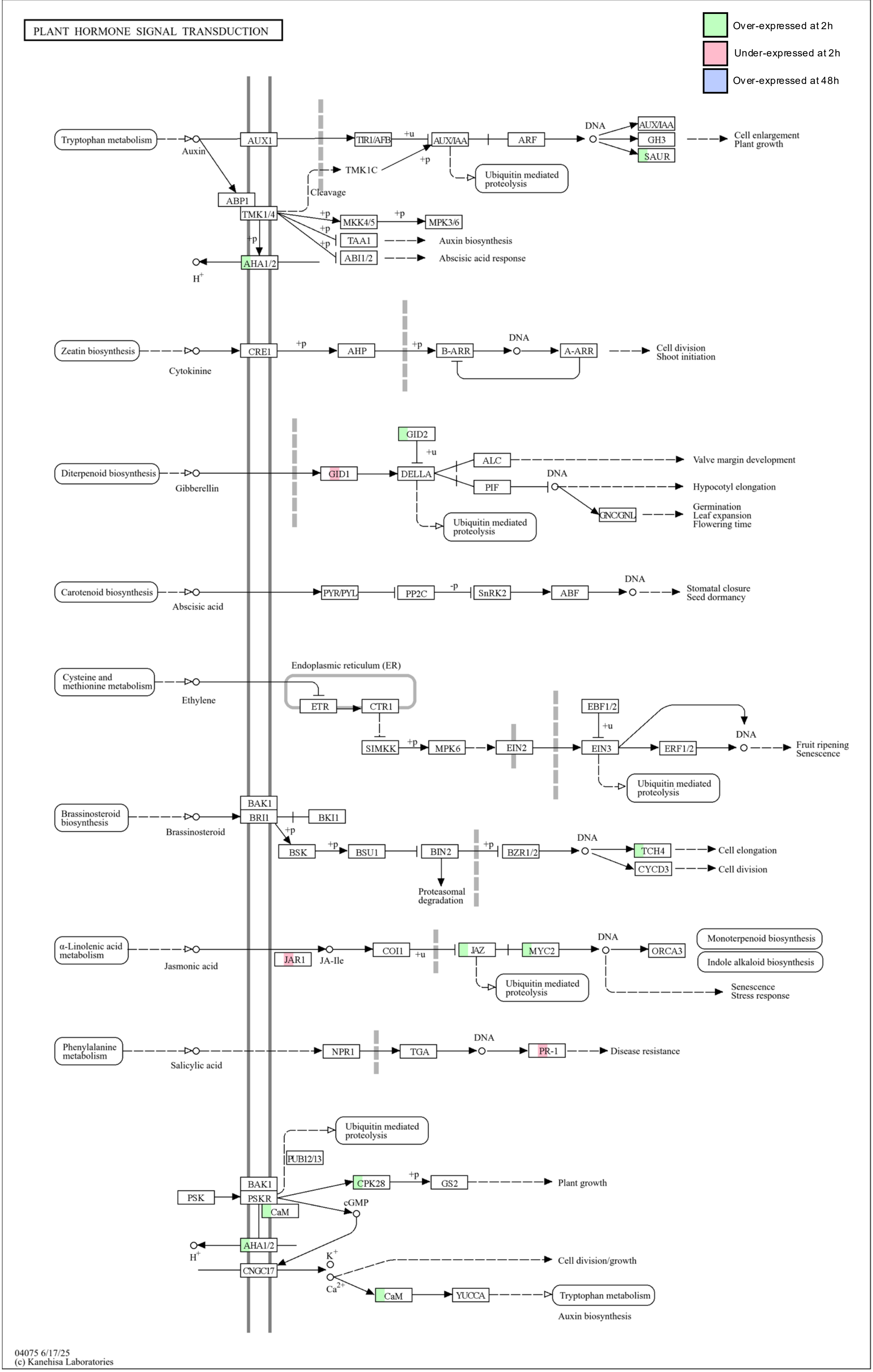

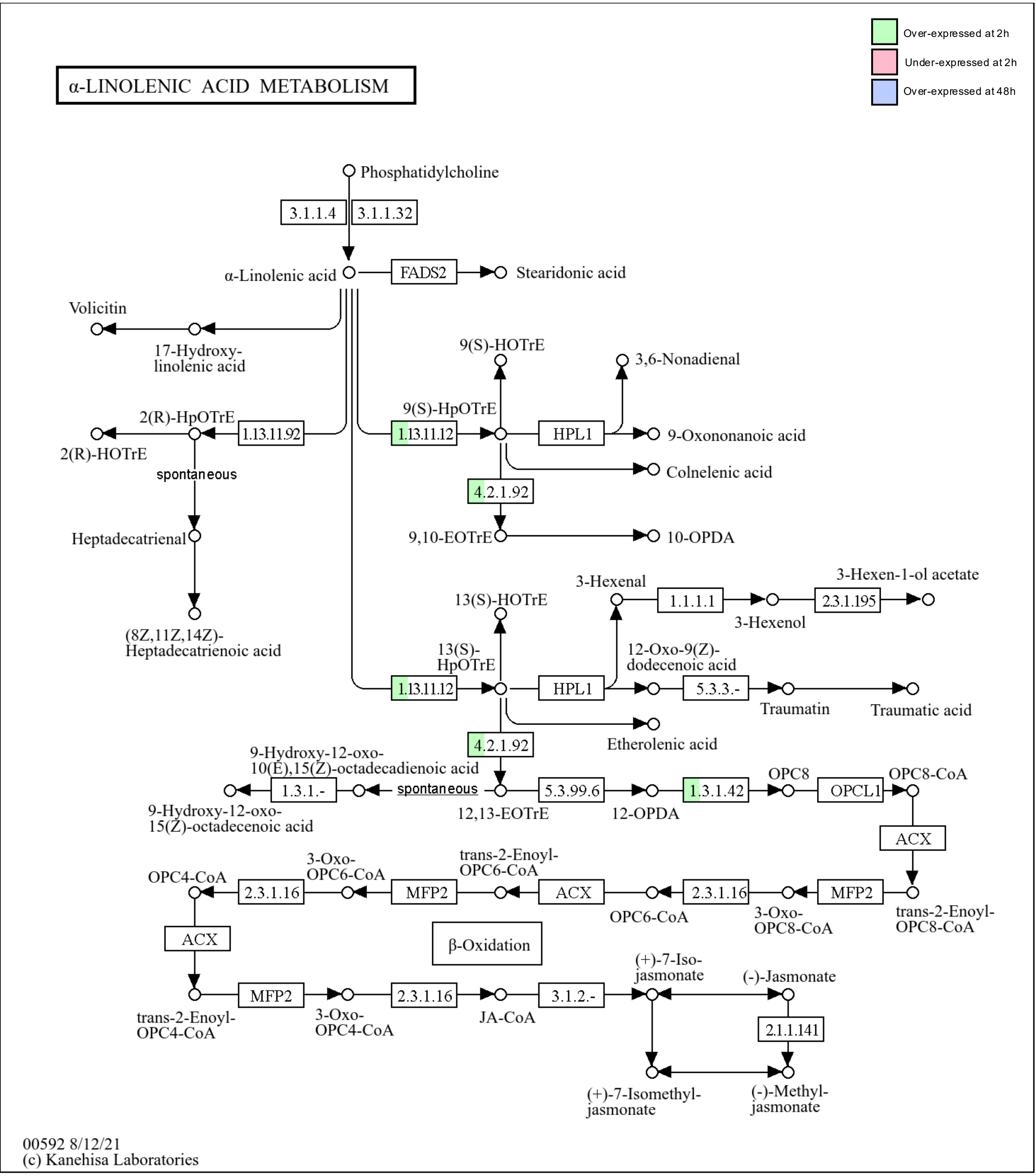

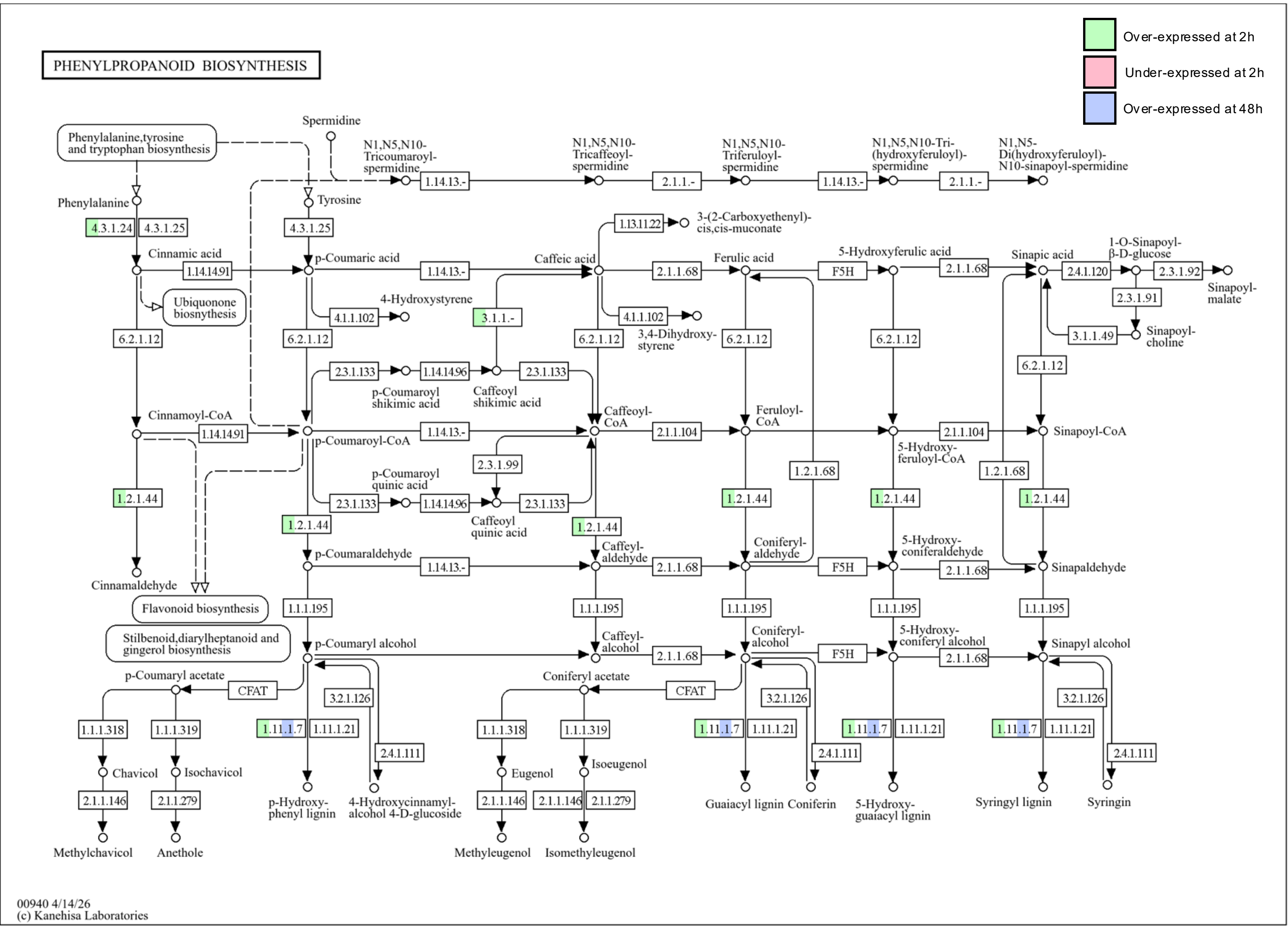

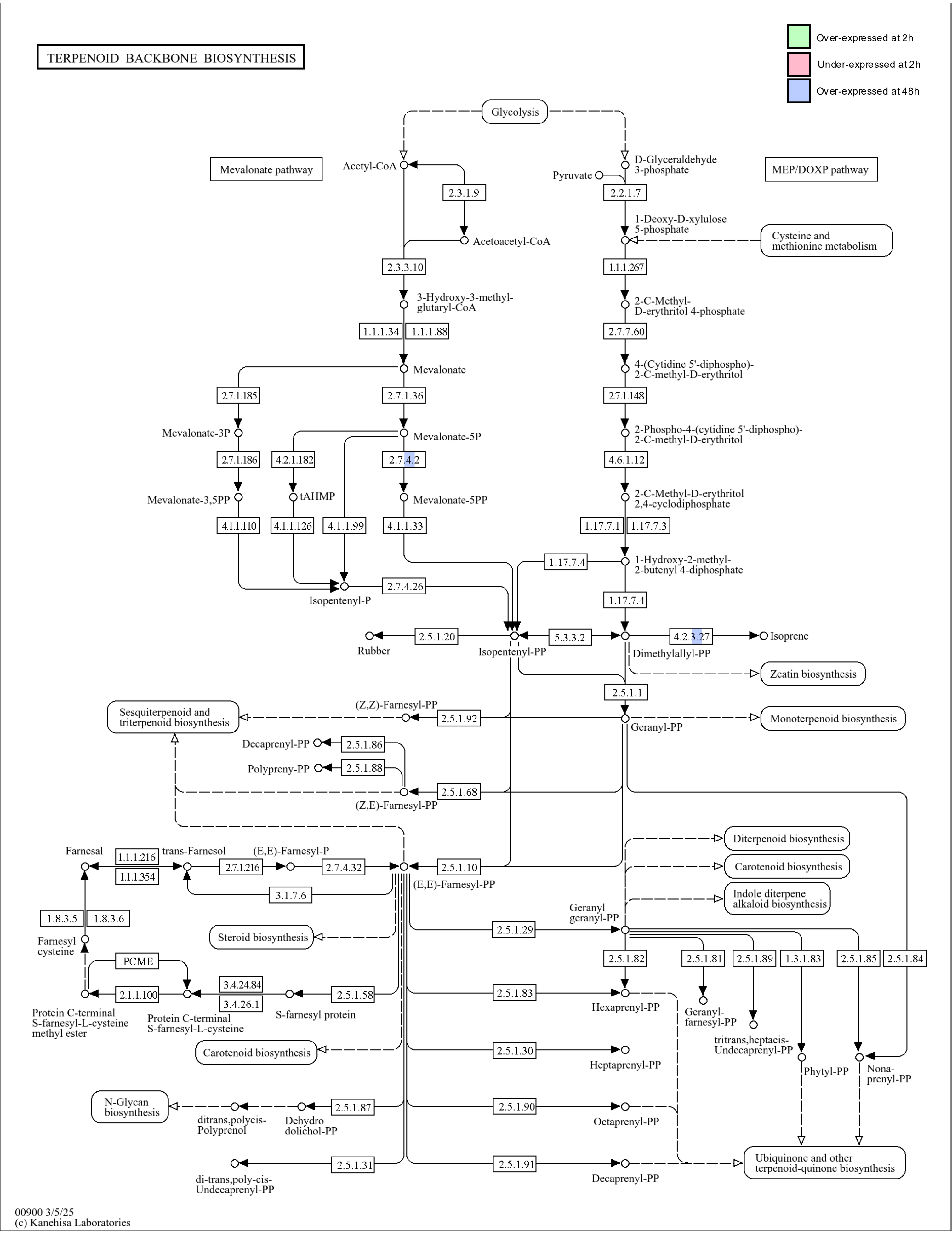

MA_107643g0010

G

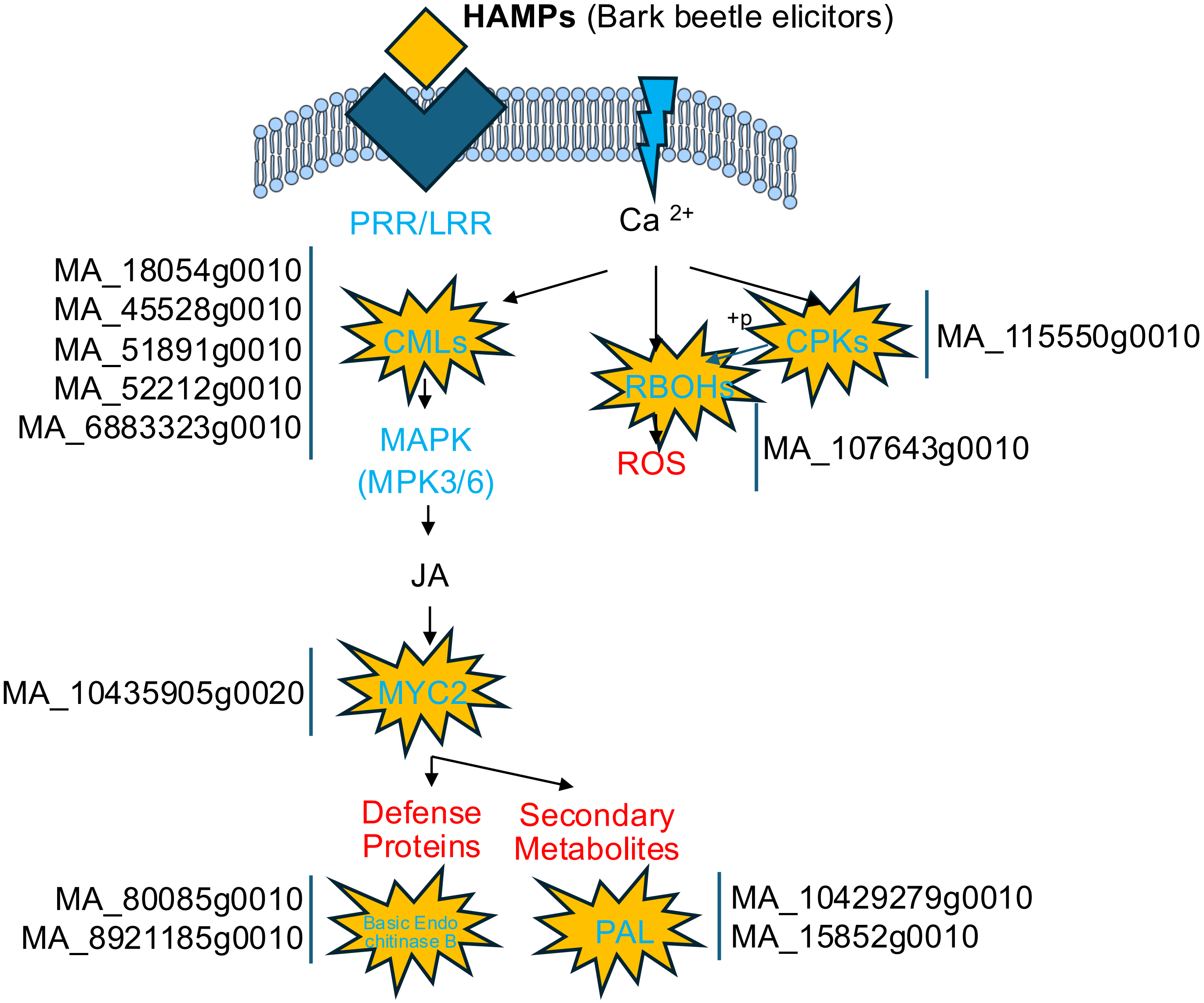

### Supplementary Figure S3

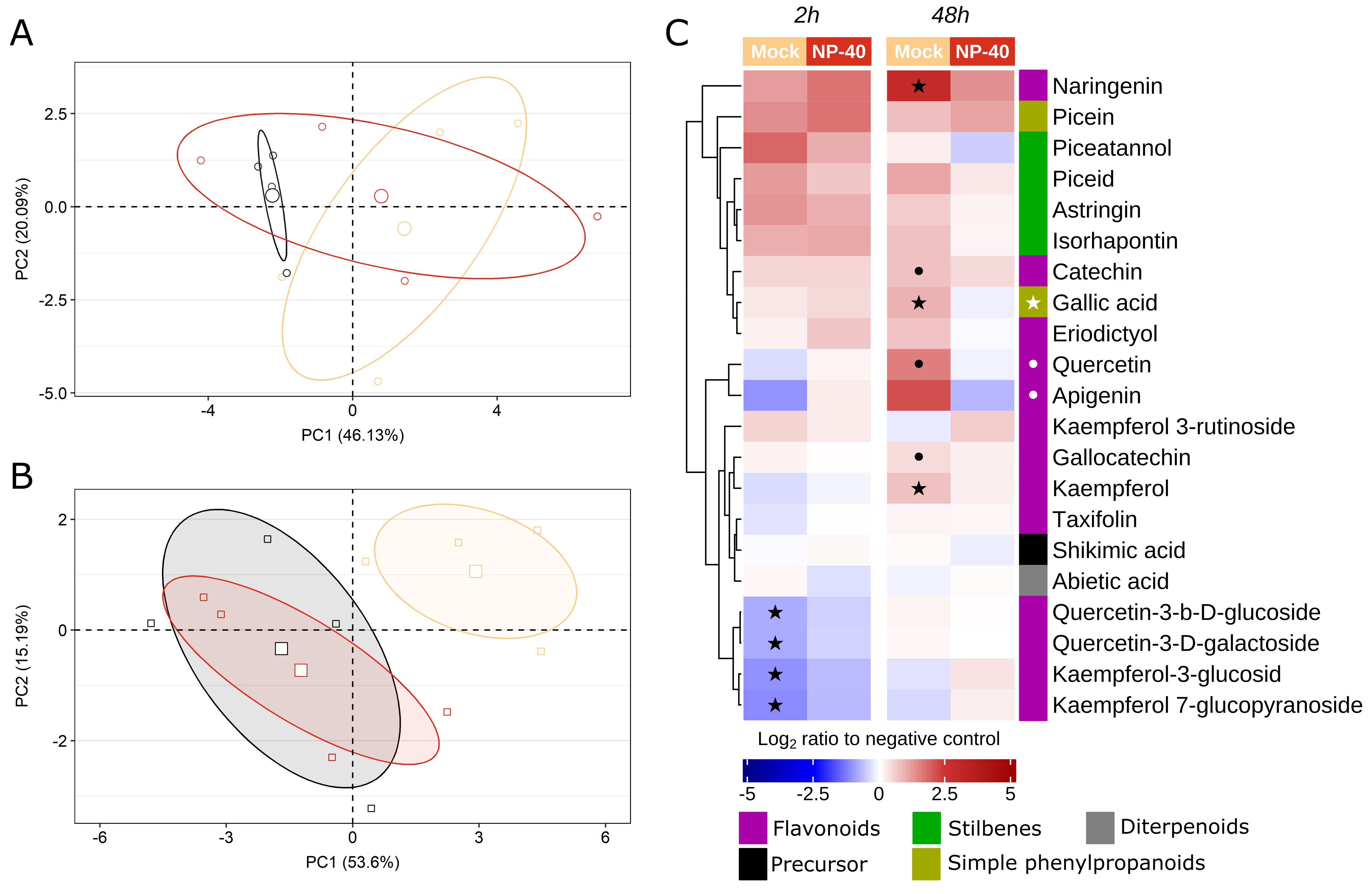
